## Supplemental material for "The G protein alpha Chaperone and Guanine-Nucleotide Exchange Factor RIC-8 Regulates Cilia Morphogenesis in *Caenorhabditis elegans* Sensory Neurons"

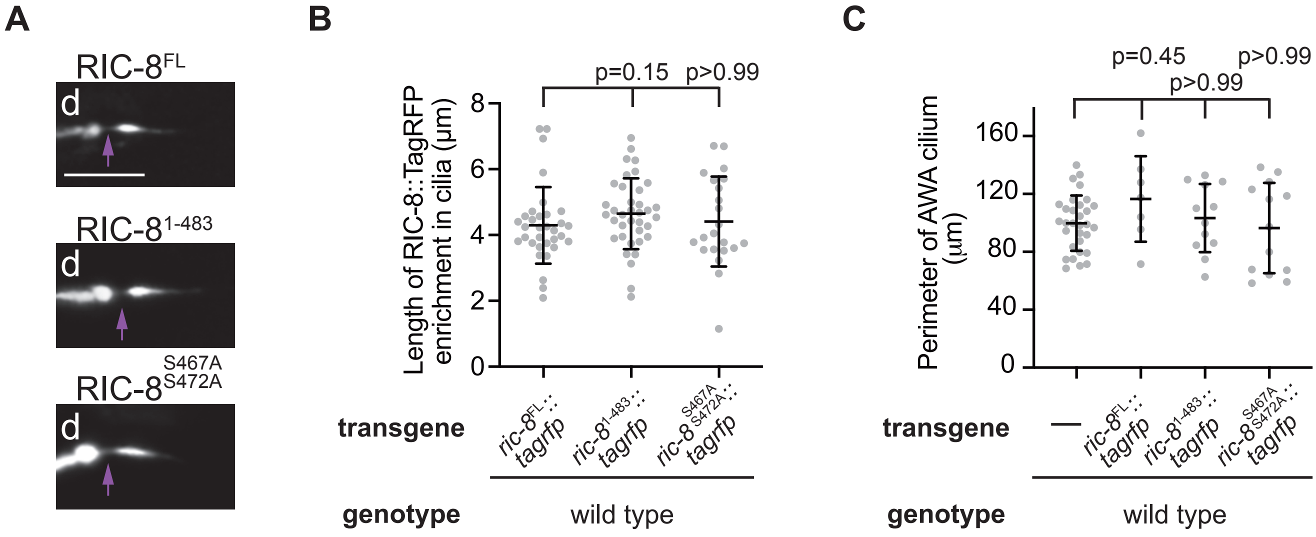
A
RIC-8FL
d
RIC-81-483
d
RIC-8S467A S472A
d
B
p=0.15
p>0.99
Length of RIC-8::TagRFP enrichment in cilia (µm)
8
6
4
2
0
transgene
ric-8FL::tagrfp
ric-81-483::tagrfp
ric-8S467A S472A::tagrfp
genotype
wild type
C
p=0.45
p>0.99
p>0.99
Perimeter of AWA cilium (µm)
160
120
80
40
0
transgene
—
ric-8FL::tagrfp
ric-81-483::tagrfp
ric-8S467A S472A::tagrfp
genotype
wild type

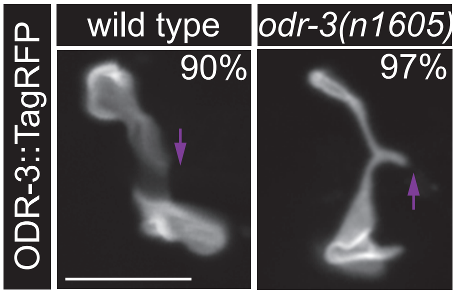


Supplemental Figure 2

Campagna *et al*.

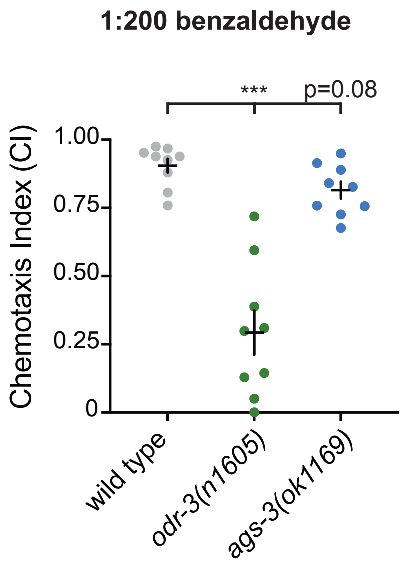
1:200 benzaldehyde
***
p=0.08
Chemotaxis Index (CI)
1.00
0.75
0.50
0.25
0
wild type
*odr-3(n1605)*
*ags-3(ok1169)*

**Supplemental Table 1:** List of *C. elegans* strains used in this work

| Strain | Genotype | Source |
|---|---|---|
| PY1089 | *kyIs104[str-1*p*::gfp]* X | (1) |
| NWM017 | *ric-8(md303)* IV; *kyIs104[str-1*p*::gfp]* X | This work |
| NWM272 | *ric-8(md1909)* IV; *kyIs104[str-1*p*::gfp]* X | This work |
| NWM089 | *ric-8(ok98)/nT1[qIs51]* IV; *kyIs104[str-1*p*::gfp]* X | This work |
| PSAB1120 | *oyIs88[gpa-4Δ6*p*::myr-gfp]* | (2) |
| PY3453 | *oyIs50[ceh-36*p*::gfp]* IV | (3) |
| NWM058 | *ric-8(ok98)/nT1[qIs51]* IV; *oyIs88[gpa-4Δ6*p*::myr-gfp]* | This work |
| NWM059 | *ric-8(ok98)/nT1[qIs51]* IV; *nchEx015[odr-1*p*::rfp]* | This work |
| NWM237 | *ric-8(md1909)* IV; *oyIs50[ceh-36*p*::gfp]* IV | This work |
| NWM386 | *ric-8(md1909)* IV; *oyIs50[ceh-36*p*::gfp]* IV; *nchEx001[ceh-36Δ*p*::ric-8::tagrfp, unc-122Δ*p*::dsRed]* | This work |
| NWM169 | *ric-8(md303)* IV; *nuIs11[osm-10*p*::gfp + lin-15(+)]* | This work |
| HA3 | *nuIs11[osm-10*p*::gfp + lin-15(+)]* | CGC |
| NWM108 | *nchEx002[bbs-8*p*::myrgfp, bbs-8*p*::ric-8::tagrfp, unc-122Δ*p*::gfp]* | This work |
| NWM393 | *nchEx003[bbs-8*p*::ric-8::tagrfp, bbs-8*p*::nphp-2s::gfp, unc-122Δ*p*::dsRed]* | This work |
| NWM344 | *oyIs50[ceh-36*p*::gfp]* IV; *nchEx001[ceh-36Δ*p*::ric-8::tagrfp, unc-122Δ*p*::dsRed]* | This work |
| NWM031 | *nchEx004[bbs-8*p*::ric-8::tagrfp, unc-122Δ*p*::gfp]*; *nchEx013[nphp-4*p*::nphp-4::gfp, unc-122Δ*p*::dsRed]* | This work |
| NWM036 | *ric-8(md303)* IV; *oyIs88[gpa-4Δ6*p*::myr-gfp]* | This work |
| NWM133 | *ric-8(md303)* IV; *oyIs88[gpa-4Δ6*p*::myr-gfp]*; *nchEx004[bbs-8*p*::ric-8::tagrfp, unc-122Δ*p*::gfp]* | This work |
| NWM187 | *ric-8(md303)* IV; *oyIs88[gpa-4Δ6*p*::myr-gfp]*; *nchEx005[bbs-8*p*::ric-8*$^{1-483}$*::tagrfp, unc-122Δ*p*::gfp]* | This work |
| NWM260 | *ric-8(md1909)* IV; *oyIs88[gpa-4Δ6*p*::myr-gfp]*; *nchEx006[bbs-8*p*::ric-8*$^{1-522}$*::tagrfp, unc-122Δ*p*::gfp]* | This work |
| NWM236 | *ric-8(md303)* IV; *oyIs88[gpa-4Δ6*p*::myr-gfp]*; *nchEx007[bbs-8*p*::ric-8*$^{S467A,S472A}$*::tagrfp, unc-122Δ*p*::gfp]* | This work |
| NWM278 | *oyIs88[gpa-4Δ6*p*::myr-gfp]*; *nchEx004[bbs-8*p*::ric-8::tagrfp, unc-122Δ*p*::gfp]* | This work |
| NWM219 | *oyIs88[gpa-4Δ6*p*::myr-gfp]*; *nchEx005[bbs-8*p*::ric-8*$^{1-483}$*::tagrfp, unc-122Δ*p*::gfp]* | This work |
| NWM226 | *oyIs88[gpa-4Δ6*p*::myr-gfp]*; *nchEx007[bbs-8*p*::ric-8*$^{S467A,S472A}$*::tagrfp, unc-122Δ*p*::gfp]* | This work |
| NWM185 | *oyIs50[ceh-36*p*::gfp]* IV; *odr-3(n1605)* V | This work |
| NWM286 | *oyIs50[ceh-36*p*::gfp]* IV; *ric-8(md1909)* IV; *odr-3(n1605)* V | This work |
| NWM313/NWM390 | *oyIs50[ceh-36*p*::gfp]* IV; *nchEx008[ceh-36Δ*p*::odr-3::tagrfp, unc-122Δ*p*::gfp]* | This work |
| NWM262/NWM243 | *oyIs50[ceh-36*p*::gfp]* IV; *odr-3(n1605)* V; *nchEx008[ceh-36Δ*p*::odr-3::tagrfp, unc-122Δ*p*::gfp]* | This work |
| NWM292 | *nchEx009[hsp-16.2*p*::ric-8::vc155, ceh-36Δ*p*::odr-3::vn173, unc-122Δ*p*::dsRed]* | This work |

| | | |
|---|---|---|
| NWM273 | *nchEx010[hsp-16.2*p*::vc155, ceh-36Δ*p*::vn173, unc-122Δ*p*::dsRed]* | This work |
| NWM412 | *nchEx011[hsp-16.2*p*::ric-8*$^{1-483}$*::vc155, ceh-36Δ*p*::odr-3::vn173, unc-122Δ*p*::dsRed]* | This work |
| NWM413 | *oyIs50[ceh-36*p*::gfp]* IV; *ric-8(md1909)* IV; *nchEx008[ceh-36Δ*p*::odr-3::tagrfp, unc-122Δ*p*::gfp]* | This work |
| NWM397 | *nchEx012[ceh-36Δ*p*::ric-8::gfp, unc-122Δ*p*::dsRed]* | This work |
| NWM430 | *odr-3(n1605)* V; *nchEx012[ceh-36Δ*p*::ric-8::gfp, unc-122Δ*p*::dsRed]* | This work |
| NWM302 | *oyIs50[ceh-36*p*::gfp]* IV; *ags-3(ok1169)* X | This work |
| NWM396 | *oyIs50[ceh-36*p*::gfp]* IV; *ric-8(md1909)* IV; *ags-3(ok1169)* X | This work |
| NWM429 | *oyIs50[ceh-36*p*::gfp]* IV; *ags-3(ok1169)* X; *nchEx008[ceh-36Δ*p*::odr-3::tagrfp, unc-122Δ*p*::dsRed]* | This work |
| NWM452 | *ric-8(md1909)* IV; *oyIs50[ceh-36*p*::gfp]* IV; *ags-3(ok1169)* X; *nchEx008[ceh-36Δ*p*::odr-3::tagrfp, unc-122Δ*p*::dsRed]* | This work |
| N2 | *C. elegans* wild isolate | CGC |
| CX3222 | *odr-3(n1605)* V | CGC |
| RB1145 | *ags-3(ok1169)* X | CGC |
| NWM431 | *oyIs50[ceh-36*p*::gfp]* IV; *nchEx014[ceh-36Δ*p*::grk-2*$^{CT}$*, unc-122Δ*p*::dsRed]* | This work |
| NWM453 | *ric-8(md1909)* IV; *oyIs50[ceh-36*p*::gfp]* IV; *nchEx014[ceh-36Δ*p*::grk-2*$^{CT}$*, unc-122Δ*p*::dsRed]* | This work |
| NWM352 | *oyIs50[ceh-36*p*::gfp] IV; nchEx016[ceh-36Δ*p*::odr-3*$^{Q206L}$*, unc-122Δ*p*::dsRed]* | This work |

**Supplemental Table 2:** List of plasmids used in this work

| Plasmid | Description | Source |
|---|---|---|
| NWM017 | *ceh-36Δ*p*::ric-8::tagrfp* | This work |
| Co-injection marker | *unc-122Δ*p*::dsRed* | (4) |
| NWM016 | *bbs-8*p*::myr-gfp* | This work |
| NWM005 | *bbs-8*p*::ric-8::tagrfp* | This work |
| Co-injection marker | *unc-122Δ*p*::gfp* | (4) |
| NWM047 | *bbs-8*p*::nphp-2s::gfp* | This work |
| NWM010 | *bbs-8*p*::ric-8*$^{1\text{-}483}$*::tagrfp* | This work |
| NWM029 | *bbs-8*p*::ric-8*$^{1\text{-}522}$*::tagrfp* | This work |
| NWM007 | *bbs-8*p*::ric-8*$^{S467A,S472A}$*::tagrfp* | This work |
| NWM032 | *ceh-36Δ*p*::odr-3::tagrfp* | This work |
| NWM030 | *hsp-16.2*p*::ric-8::vc155* | This work |
| NWM034 | *ceh-36Δ*p*::odr-3::vn173* | This work |
| NWM031 | *hsp-16.2*p*::vc155* | This work |
| NWM033 | *ceh-36Δ*p*::vn173* | This work |
| NWM046 | *hsp-16.2*p*::ric-8*$^{1\text{-}483}$*::vc155* | This work |
| NWM043 | *ceh-36Δ*p*::ric-8::gfp* | This work |
| NWM048 | *ceh-36Δ*p*::grk-2*$^{CT}$ | This work |
| NWM040 | *ceh-36Δ*p*::odr-3*$^{Q206L}$*::tagrfp* | This work |
